# Longitudinal characterisation of phagocytic and neutralisation functions of anti-Spike antibodies in plasma of patients after SARS-CoV-2 infection

**DOI:** 10.1101/2021.12.21.473774

**Authors:** Anurag Adhikari, Arunasingam Abayasingam, Chaturaka Rodrigo, David Agapiou, Elvis Pandzic, Nicholas A Brasher, Bentotage Samitha Madushan Fernando, Elizabeth Keoshkerian, Hui Li, Ha Na Kim, Megan Lord, Gordona Popovic, William Rawlinson, Michael Mina, Jeffrey J Post, Bernard Hudson, Nicole Gilroy, Adam W. Bartlett, Golo Ahlenstiel, Branka Grubor-Bauk, Dominic Dwyer, Pamela Konecny, Andrew R Lloyd, Marianne Martinello, Rowena A Bull, Nicodemus Tedla, on behalf of the COSIN study group

**Affiliations:** School of Medical Sciences, Faculty of Medicine, UNSW Australia, Sydney, NSW, Australia; The Kirby Institute, UNSW Australia, Sydney, NSW, Australia; Department of Infection and Immunology, Kathmandu Research Institute for Biological Sciences, Lalitpur, Nepal; Katharina Gaus Light Microscopy Facility, Mark Wainwright Analytical Centre, University of New South Wales, Sydney, NSW, Australia; School of Biomedical Engineering, Faculty of Engineering, UNSW Australia, Sydney, NSW, Australia; School of Mathematics and Statistics, University of New South Wales, Sydney, NSW, Australia; Serology and Virology Division, Department of Microbiology, NSW Health Pathology, Prince of Wales Hospital, Sydney, New South Wales, Australia; Northern Beaches Hospital, NSW, Australia; Prince of Wales Clinical School, UNSW Australia, Sydney, NSW Australia; Royal North Shore Hospital, Sydney, New South Wales, Australia; Westmead Hospital, Sydney, New South Wales, Australia; Blacktown Mt Druitt Hospital, Blacktown, NSW, Australia; Viral Immunology Group, Adelaide Medical School, University of Adelaide and Basil Hetzel Institute for Translational Health Research, Adelaide, SA, Australia; Institute of Clinical Pathology and Medical Research, Westmead Hospital, NSW, Australia; St. George Clinical School, University of New South Wales, NSW, Australia

## Abstract

Phagocytic responses by effector cells to antibody or complement-opsonised viruses have been recognized to play a key role in anti-viral immunity. These include antibody dependent cellular phagocytosis mediated via Fc-receptors, phagocytosis mediated by classically activated complement-fixing IgM or IgG1 antibodies and antibody independent phagocytosis mediated via direct opsonisation of viruses by complement products activated via the mannose-binding lectin pathway. Limited data suggest these phagocytic responses by effector cells may contribute to the immunological and inflammatory responses in SARS-CoV-2 infection, however, their development and clinical significance remain to be fully elucidated. In this cohort of 62 patients, acutely ill individuals were shown to mount phagocytic responses to autologous plasma-opsonised SARS-CoV-2 Spike protein-coated microbeads as early as 10 days post symptom onset. Heat inactivation of the plasma prior to use as an opsonin caused 77-95% abrogation of the phagocytic response, and pre-blocking of Fc-receptors on the effector cells showed only 18-60% inhibition. These results suggest that SARS-CoV-2 can provoke early phagocytosis, which is primarily driven by heat labile components, likely activated complements, with variable contribution from anti-Spike antibodies. During convalescence, phagocytic responses correlated significantly with anti-Spike IgG titers. Older patients and patients with severe disease had significantly higher phagocytosis and neutralisation functions when compared to younger patients or patients with asymptomatic, mild, or moderate disease. A longitudinal study of a subset of these patients over 12 months showed preservation of phagocytic and neutralisation functions in all patients, despite a drop in the endpoint antibody titers by more than 90%. Interestingly, surface plasmon resonance showed a significant increase in the affinity of the anti-Spike antibodies over time correlating with the maintenance of both the phagocytic and neutralisation functions suggesting that improvement in the antibody quality over the 12 months contributed to the retention of effector functions.

**Author Summary:** Limited data suggest antibody dependent effector functions including phagocytosis may contribute to the immunological and inflammatory responses in SARS CoV-2 infection, however, their development, maintenance, and clinical significance remain unknown. In this study we show:

1. Patients with acute SARS CoV-2 infection can mount phagocytic responses as early as 10 days post symptom onset and these responses were primarily driven by heat labile components of the autologous plasma. These results indicate that the current approach of studying phagocytosis using purified or monoclonal antibodies does not recapitulate contribution by all components in the plasma.
2. In convalescent patients, high phagocytic responses significantly correlated with increasing age, increasing disease severity, high neutralisation functions and high anti-Spike antibody titers, particularly IgG1.
3. Longitudinal study of convalescent patients over a 12-month period showed maintenance of phagocytic and neutralisation functions, despite a drop in the anti-Spike endpoint antibody titers by more than 90%. However, we found significant increase in the affinity of the anti-Spike antibodies over the 12-month period and these correlated with the maintenance of functions suggesting that improvement in the antibody quality over time contributed to the retention of effector functions. Clinically, measuring antibody titers in sera but not the quality of antibodies is considered a gold standard indicator of immune protection following SARS-CoV 2 infection or vaccination. Our results challenge this notion and recommends change in the current clinical practice.

## Introduction

Since emerging in 2019, SARS-CoV-2 has spread rapidly worldwide infecting over 265 million individuals with over five million deaths [1]. To understand disease pathogenesis and immune correlates of protection, detailed characterisation of the immune response against SARS-CoV-2 both during the acute phase and longitudinally is required including in patients varying in age and disease severity.

Early polyfunctional T cell, B cell and antibody responses against SARS-CoV-2 are associated with reduced clinical severity and better disease outcomes [2]. At the beginning of the pandemic there was much controversy over the protective capacity of antibodies in coronavirus infections [3,4]. In rodent models, neutralising antibodies were demonstrated to block infection of SARS-CoV [5] and SARS-CoV-2 [6]. However, neutralisation alone may be insufficient for protection [7], whereas the combination of neutralising antibodies with Fc receptor dependent effector functions may reduce disease severity [8,9].

Limited data suggest that phagocytic responses by effector cells to antibody- or complement-opsonised viruses may play a key role in the immunological [9,10] and inflammatory responses to SARS-CoV-2 infection [11–14]. These include the association of antibody dependent cellular phagocytosis (ADCP) with protection [10], correlation of earlier IgG class switching and maintenance of Fc receptor binding properties with reduction in disease severity [9]. By contrast, phagocytosis mediated by classically activated complement-fixing IgM antibodies has been associated with unfavourable clinical outcomes [15], and antibody independent phagocytosis mediated by complement products activated via the mannose-binding lectin pathway were associated with a hyperinflammatory state [11]. However, the kinetics of development of these phagocytic responses, their maintenance, and their clinical significance remain to be fully elucidated. In particular, little is known about the phagocytic responses during acute disease, and the long-term retention of these functions and the associated predictive factors in convalescent patients beyond five months are unknown [16]. Importantly, studies to date used purified patient antibodies or cloned monoclonal antibodies to detect phagocytosis and so disregarded potentially significant contribution of plasma proteins such as activated complements that are relevant in phagocytic responses against respiratory pathogens [11,17,18].

In this study acutely ill patients with COVID-19 mounted phagocytic responses to autologous plasma-opsonised-SARS-CoV-2 Spike protein-coated microbeads as early as 10 days post symptom onset, independent of disease severity and despite variable levels of anti-Spike antibody titers. Heat inactivation of the plasma prior to use as an opsonin caused 77-95% abrogation of the phagocytic response and blocking of the Fc-receptors revealed 18-60% inhibition. These results suggest that SARS-CoV-2 can provoke early phagocytosis which is primarily driven by heat labile components in the plasma with variable contribution from anti-Spike antibodies. In convalescent patients, there was an increase in phagocytosis which correlated significantly with higher anti-Spike IgG titers and with antibody mediated neutralisation. Older patients and patients with severe disease had significantly higher phagocytic responses when compared to younger patients and patients with asymptomatic, mild, or moderate disease. This age and disease severity dependent difference in the anti-Spike antibody dependent phagocytosis mirrored the anti-Spike neutralisation activity. There was no significant difference in phagocytic responses between men and women. Longitudinally, phagocytosis and neutralisation functions were still detectable at the 12-month timepoint, in parallel with a significant increase in affinity of the anti-Spike antibodies, despite an over 90% decline of the endpoint antibody titers. These findings suggest that improved quality of the antibodies, likely due to somatic hypermutation, may play a major role in the preservation of these effector functions.

## Results

### Study cohort

Antibody assays were performed on plasma and serum samples collected from 62 participants that included six acutely ill (two females, median age 52.5 [range: 14-79] (Table 1) and 56 convalescent patients (25 females, median age 53 [range: 19-94] (Table 1).

**Table 1:**
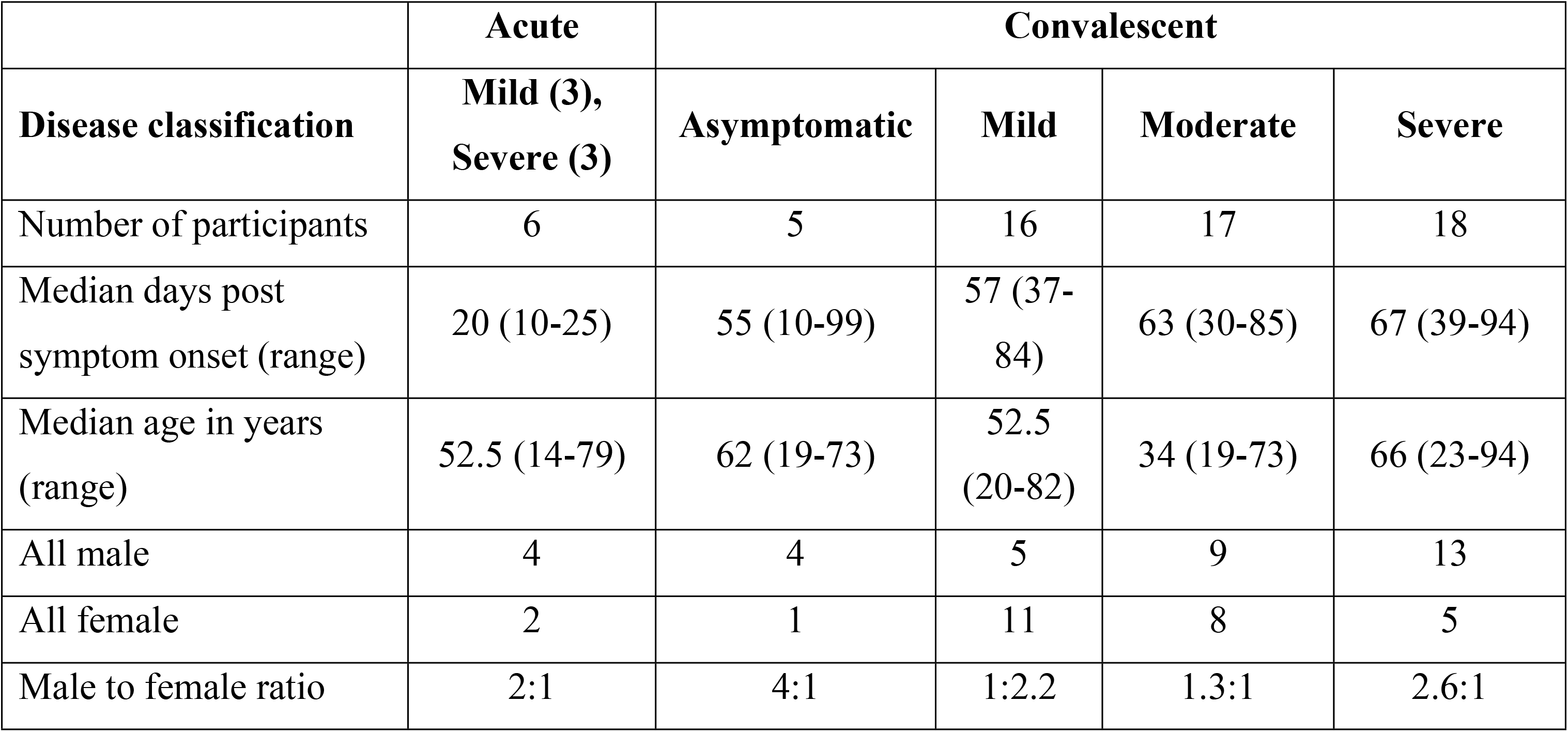
Summary of demographic profile of the patients with COVID-19.

|  | Acute | Convalescent |  |  |  |
| --- | --- | --- | --- | --- | --- |
| Disease classification | Mild (3),<br>Severe (3) | Asymptomatic | Mild | Moderate | Severe |
| Number of participants | 6 | 5 | 16 | 17 | 18 |
| Median days post symptom onset (range) | 20 (10-25) | 55 (10-99) | 57 (37-84) | 63 (30-85) | 67 (39-94) |
| Median age in years (range) | 52.5 (14-79) | 62 (19-73) | 52.5 (20-82) | 34 (19-73) | 66 (23-94) |
| All male | 4 | 4 | 5 | 9 | 13 |
| All female | 2 | 1 | 11 | 8 | 5 |
| Male to female ratio | 2:1 | 4:1 | 1:2.2 | 1.3:1 | 2.6:1 |

For the acutely ill patients, samples were collected 10-25 days post symptom onset (DPS) (median: 20 DPS) (Table 1) and for patients in convalescence, samples were collected on days 10-99 DPS (median: 59.5 DPS) (visit 1). In a subset of the convalescent patients (n=9), longitudinal samples were collected on 110-252 DPS (median: 150 DPS) (visit 2) and on 329-381 DPS (median: 345 DPS) (visit 3), respectively. Among the convalescent patients, five were asymptomatic, 16 had mild, 17 had moderate and 18 had severe disease, and all groups had comparable median DPS (Table 1). As expected, the median age for the patients with severe disease was higher (66 years) while for those with moderate disease was much lower (34 years) than the overall median age of 53 years. The male to female ratio (1.2:1) of the cohort was representative of the epidemic, however in the severe disease group males were over-represented at a ratio of 2.6 to 1 and under-represented in patients with mild disease in a ratio of 1 to 2.2 (Supplementary Table 1). At the time of collection of the 12-month samples, Australia had limited community transmission following the first wave in 2020 when these participants were infected [19]. Although the specific SARS-CoV-2 variants in these patients were not genotyped, all predate infections with the Delta and Omicron variants. None of these participants had received a SARS-CoV-2 vaccine at the time of sampling. A detailed description of the patients is shown in Supplementary Table 1.

### Early detection of heat labile phagocytosis in acutely ill patients

Plasma from all six acutely ill participants demonstrated detectable phagocytosis of microbeads coated with SARS-CoV-2 Spike protein as early as 10 days post symptom onset regardless of disease severity, and despite 2/6 having no detectable anti-Spike antibody end point titers (EPTs, see methods?) (Fig 1A). The gating strategy used to determine the proportion of positive cells and their mean fluorescence intensity (MFI), as well as microscopic validation for intracellular uptake of the opsonised microbeads is presented in Supplementary Fig 1. In the patients (n = 4) with detectable anti-Spike antibodies, the mean EPTs was 11015, 95% CI [−6490, 28500] with mean Spike phagocytosis score (p-score, see Methods) of 51.4, 95% CI [18.2, 84.6]. For the 2/6 patients with undetectable EPTs, the Spike p-scores were markedly lower (2.38, 95% CI [1.15, 3.61]), whereas the background cut-off in samples from unexposed healthy control subjects was 0.9, 95% CI [0.518, 1.28] (Fig 1A). Although the number of the acutely ill patients is limited, those with high Spike p-scores had low viral loads, whereas individuals with low p-scores had relatively high viral loads (Fig 1B). However, there was no correlation between the anti-Spike EPTs and Spike p-scores (Spearman’s r = 0.8, p = 0.07, Fig 1C), suggesting contribution from other components of the plasma including the heat labile complements. To confirm this, plasma from these patients was heat inactivated prior to use as an opsonin and found a profound abrogation of phagocytosis by 77-95% in 5/6 of patients (mean Spike p-score 5.7, 95% CI [−5.5, 16.9]) (Fig 1D). By contrast, Fc receptor blocking experiments indicated that anti-Spike antibodies in the plasma of these patients contributed to 18-60% of the phagocytic function (Fig 1E). Interestingly, in one patient (Patient #1) with the undetectable anti-Spike antibodies (Fig 1A), blocking of the Fc receptor showed 60.3% inhibition of phagocytosis suggesting the presence of small amounts of effective anti-Spike antibodies that were not detected by endpoint titration in ELISA. To test this further, binding assay using surface plasmon resonance were performed and revealed extremely high affinity binding of the plasma to recombinant Spike protein (KD = 4×10^−12^M) (Fig 1F) supporting the notion of low titre high affinity antibodies.

**Fig 1.**
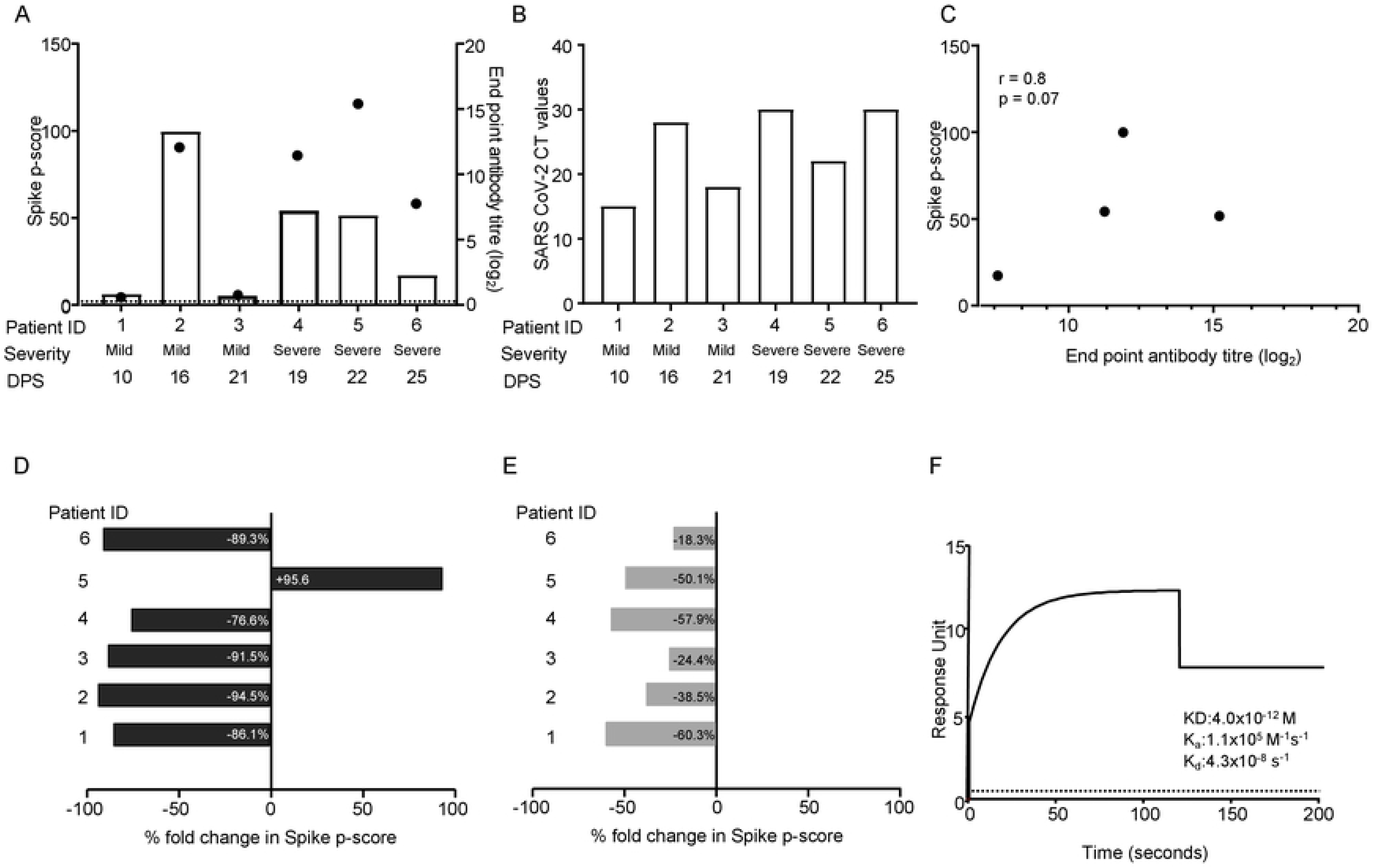
Detection of heat labile Spike phagocytosis in acutely ill patients with COVID-19. (A) A histogram showing positive Spike phagocytosis scores was detected in all acutely sick patients despite varying anti-Spike antibody EPTs (black dots), differences in DPS onset and disease severity; positive phagocytosis is defined as three standard deviations above the mean a phagocytosis scores of healthy controls (dotted line). (B) Viral load in plasma at the time of sample collection (DPS) among the patients. (C) Spearman correlation study showing a trend of positive correlation between Spike antibody EPTs and Spike p-scores despite the small sample size (n = 6, r = 0.8, p = 0.07). (D) Heat inactivation of the plasma caused a profound abrogation of Spike p-score by 77-95% in 5/6 of the patients with acute infection (mean Spike p-score 5.7, 95% CI [−5.5, 16.9]), suggesting major contribution by heat labile components of the plasma. (E) Fc receptor blocking experiments with untreated plasma indicated that the antibodies contributed to 18-60% (mean Spike p-score 17.82, 95% CI [−0.58, 36.2]) of the phagocytic function all acute patients. (F) Surface plasmon resonance showing high affinity binding of plasma obtained from patient #1 to recombinant Spike protein (KD = 4×10^−12^M) despite having undetectable anti-Spike antibody (A) and minimal Fc-receptor dependent phagocytosis (E).

### Disease severity dependent differences in phagocytosis, endpoint antibody titers and neutralisation function at convalescence

Measurement of anti-SARS-CoV-2 antibody EPT in sera collected during convalescence in the 10-99 DPS window (n=56) showed that 100% had detectable anti-Spike IgG and 98% had anti-RBD IgG antibodies (Fig 2A-B). Further analysis of the IgG subclasses of the anti-Spike responses showed that the antibodies were primarily IgG1 and IgG3 subtypes, but not IgG2 or IgG4 (Fig 2C). There was also variable levels of anti-Spike IgA and IgM antibodies (Fig 2C). Functionally, 95% (53/56) of all patient samples demonstrated significant phagocytosis of Spike protein-coated microbeads (Fig 2D), 84% (47/56) had phagocytosis of RBD protein-coated microbeads, albeit at lower phagocytic scores (Fig 2E), and 86% (48/56) could neutralise Spike pseudovirus (Fig 2F).

**Fig 2:**
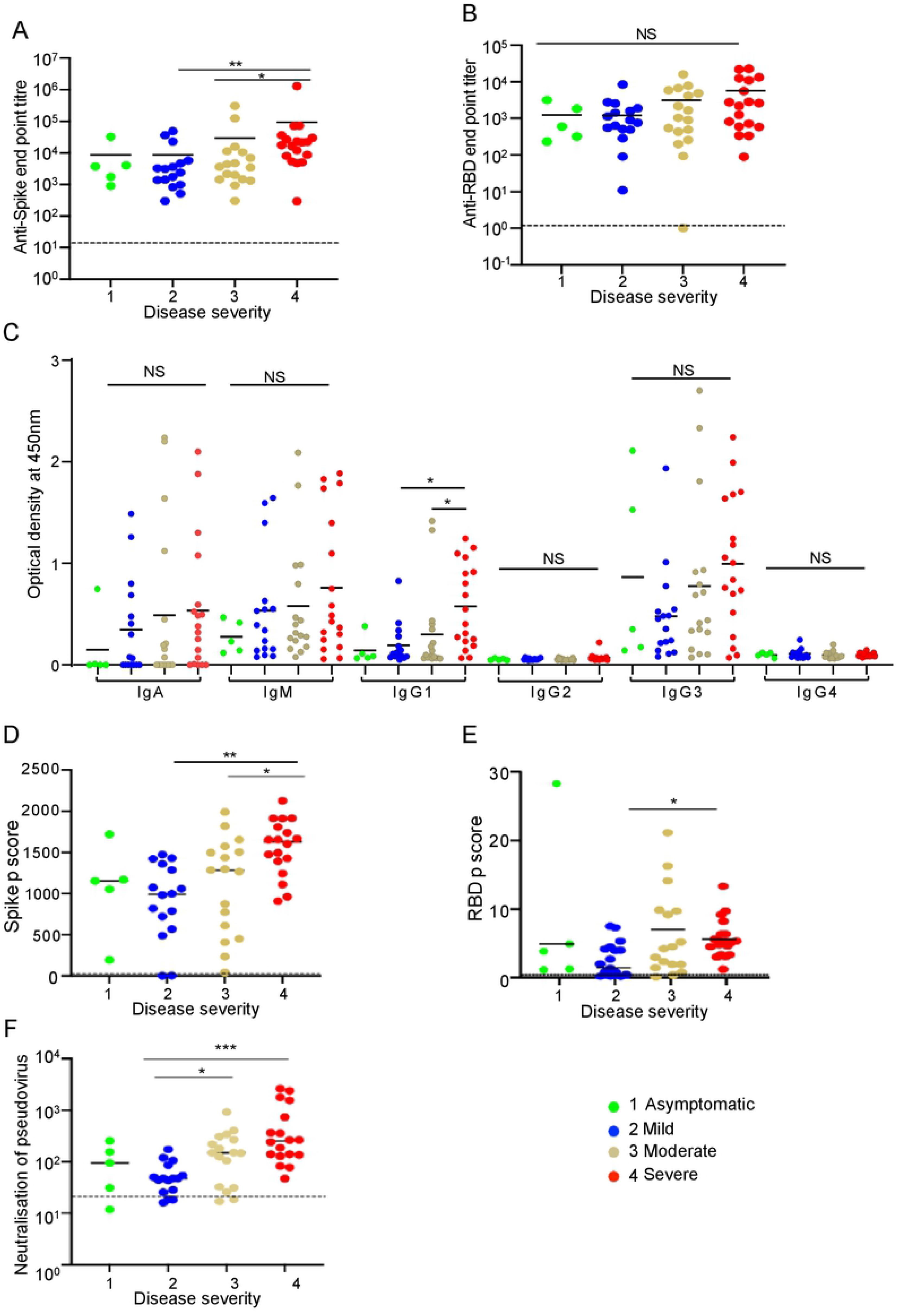
Disease severity dependent differences in endpoint antibody titers, phagocytosis and neutralisation titre at convalescence. (A) Disease severity dependent increase in anti-Spike antibody EPTs in plasma of patients was significant in patients with severe disease when compared to moderate (difference of mean = 64498.1, 95% CI [−7340, 66700]), mild (difference of mean = 85558.5, 95% CI [1550, 15600]); B) Anti-RBD antibody EPTs showing high level expression in all patients but there was no significant difference among the various patient groups. (C) Analysis of the anti-Spike antibody classes and IgG subtypes showed IgG1 as the major antibody subtype contributing to the significant increase in the anti-Spike EPTs in the patients with severe disease. (D) Severe disease group has significantly higher Spike p-score compared to moderate (difference of mean = −454.9, 95% CI [−885.3, −24.38]) and mild diseased group (difference of mean = −650.6, 95% CI [−1088, −213.3]. (E) Anti-RBD antibodies-mediated phagocytic response was significantly higher in the severe when compared to mild (difference of mean = 0.6, 95% CI [4.34, 7.02]) diseased individuals. (F) Severe disease group has significantly higher neutralisation titre compared to moderate (difference of mean = −440.8, 95% CI [−889.8, 8.122]) and mild diseased group (difference of mean = −586.5, 95% CI [−1043, −130.4]); F) Dotted lines in each plot show median or mean values+3SD of 25 healthy control samples (*p<0.05, **p<0.01, ***p<0.001).

Stratification by disease severity into asymptomatic (n=5), mild (n=16), moderate (n=17) or severe (n=18) disease showed a severity dependent increase in anti-Spike antibody EPTs (Figs 2A and 2C). A significant difference in anti-Spike antibody EPTs was observed in between disease severity groups (Kruskal-Wallis test, H [3] = 12.61, p = 0.005), whereas the severe group had significantly higher anti-Spike antibody EPT than the mild (Kruskal-Wallis test, p = 0.006) and moderate (Kruskal-Wallis test, p = 0.04) groups (Fig 2A). There was no significant difference in anti-RBD antibody EPT among disease severity groups (Kruskal-Wallis test, p = 0.28) (Fig 2B). Multiple comparison of the anti-Spike antibody subtypes among disease severity groups showed higher anti-Spike IgG1 (Kruskal-Wallis test, mild versus severe: p = 0.04 and moderate versus severe: p = 0.03), whereas difference between other subtypes were not statistically significant (Fig 2C). There was also a significant difference in Spike p-scores between the disease severity groups [ANOVA, F (3, 52) = 5.695, p = 0.001]. In particular, the severe disease group has significantly higher Spike p-scores compared to the moderate (mean difference = −454.9, 95% CI = −885.3 to −24.38, p = 0.03) and mild disease groups (mean difference = −650.6, 95% CI = −1088 to −213.3, p = 0.001) (Fig 2D). Similarly, a significant difference in RBD p-scores was evident between severe and mild disease groups (Kruskal-Wallis test, p = 0.04) (Fig 2E). The neutralisation titers were also significantly different between disease severity groups [ANOVA, F (3, 52) = 4.545, p = 0.006] with the severe disease group having a significantly higher neutralisation titre compared to the moderate (mean difference = −440.8, 95% CI = −889.8 to 8.122, p = 0.049) and mild disease groups (mean difference = −586.5, 95% CI = −1043 to −130.4, p = 0.006) (Fig 2F).

### Age dependent differences in phagocytosis, endpoint antibody titers and neutralisation function at convalescence

Analysis of the age stratified groups: >60 years (n=24), 40-60 years (n=14), <40 years (n=18) showed that anti-Spike antibody EPTs were significantly different between the age groups (Kruskal-Wallis test, H [2] = 13.6, p = 0.001). Patients >60 years of age had higher anti-Spike antibody EPTs compared to the 40–60-year-old group (Kruskal-Wallis test, p = 0.02) and the <40-year old patient group (Kruskal-Wallis test, p = 0.001) (Fig 3A). By contrast, there was no difference in the anti-RBD antibody EPT between the age groups (Fig 3B). Examination of the anti-Spike antibody classes and IgG subtypes revealed significant difference between the age groups (Kruskal-Wallis test, H [2] = 162.3, p = 0.0001). There were higher anti-Spike IgG1 in the >60-years old group when compared to the <40-year-old group (Kruskal-Wallis test, p = 0.0007), and significantly higher IgG3 in the >60 year group when compared to 40–60 year old group (Kruskal-Wallis test, p = 0.007). The anti-Spike IgG2 and IgG4 levels were not significantly different (Fig 3C) and there was no significant difference in Spike p-score or RBD p-score between the different age groups (Fig 3D-E). Interestingly, neuralisation titers were significantly different among the age groups (Kruskal-Wallis test, H [2] = 7.04, p = 0.029) with highest titers found in the >60 year old patients when compared to the <40 year old group (Kruskal-Wallis test, p = 0.042), (Fig 3F). There were no significant differences in anti-Spike IgG EPTs, anti-RBD IgG EPTs or Spike/RBD p-scores between male and female patients (Supplementary Fig 2A-C). However, neutralisation was significantly higher among males compared to females (Mann Whitney U-test, p = 0.01) (Supplementary Fig 2C).

**Fig 3:**
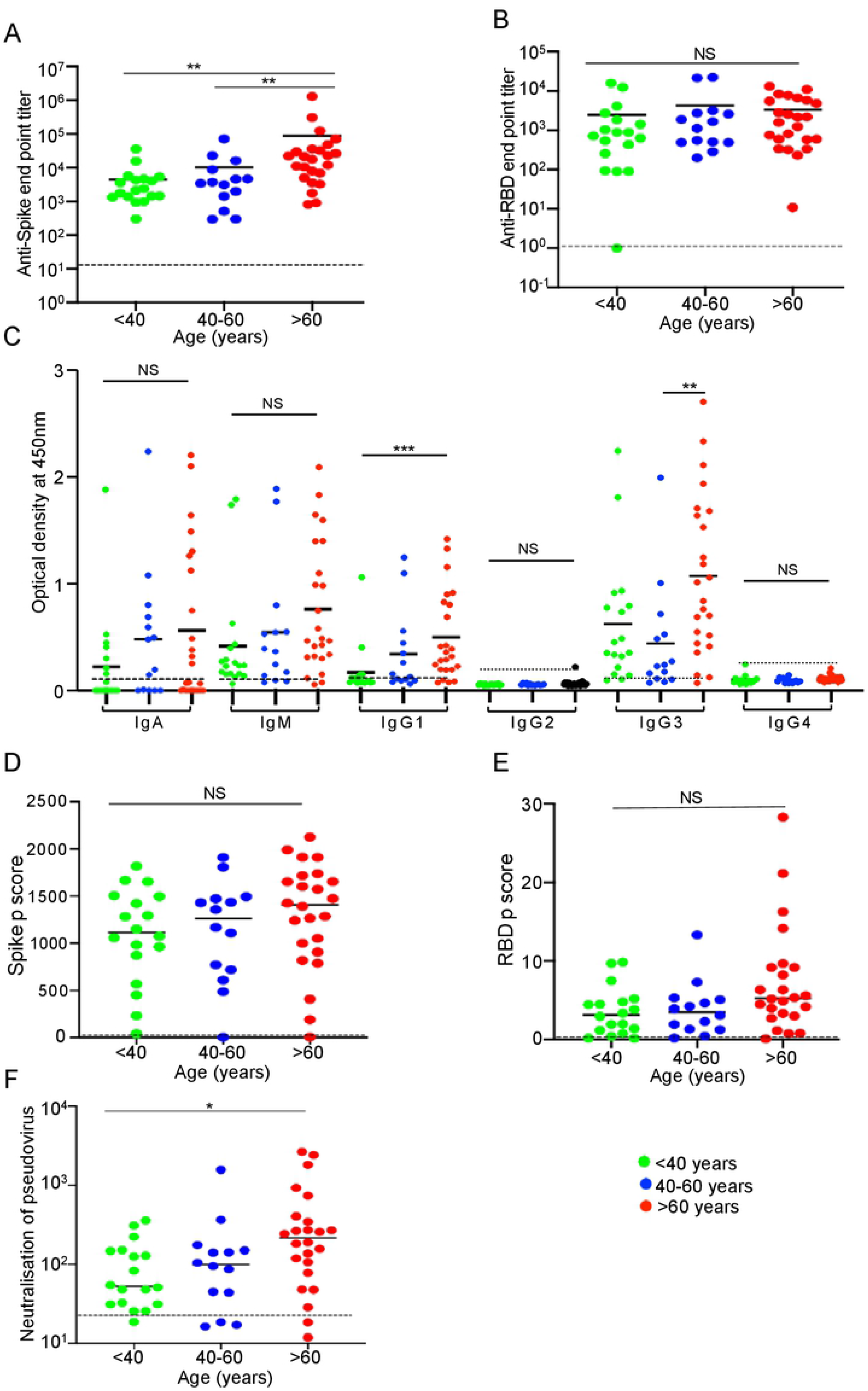
Age dependent increase in endpoint antibody titers, phagocytosis and neutralisation titre at convalescence. (A) Analysis of the age dependent responses showing the patients older than 60 years of age with higher anti-Spike antibodies EPTs than the 40-60-years old (difference of mean = 78911.2, 95% CI [390, 20200]) and <40-year-old patients (difference of mean = 83984, 95% CI [1300, 9160]). (B) However, there is no statistical difference in the anti-RBD EPT among the age groups. (C) Higher anti-Spike IgG1 among >60-years old group was found when compared to <40-years old (Kruskal-Wallis test, p = 0.0007) and significantly higher IgG3 among >60-years old group when compared to 40–60-year old (Kruskal-Wallis test, p = 0.007). (D&E) Neither the Spike p-score nor RBD p-score was statistically different among the age groups. (F) Neutralisation titre was significantly higher in the >60-year old patients when compared to the <40 (difference of mean = 382.2, 95% CI [58.4, 152]); Dotted lines in each plot show median or mean values +3SD of 25 healthy control samples (*p<0.05, **p<0.01, ***p<0.001).

### Ranking of the association of independent variables of importance to anti-Spike antibody mediated phagocytosis

Multiple linear regression analysis was performed to rank all the independent variables of importance that associated with the dependent variable (Spike p-score). Anti-Spike endpoint titre was the most important variable that predicted the Spike p-score (change in R^2^ = 0.2277) (Table 2), while the rest of the variables are listed in descending order of predictive value in Table 2. A parameter covariance matrix was plotted to determine the associative relationships between the non-categorical independent variable to yield the dependent variable Spike p-score, revealing that anti-Spike IgG1 and IgG3 associated positively with most independent variables, whereas age and DPS associated negatively or poorly with other variables (Fig 4).

**Table 2:** Variable importance ranking based on multiple regression model.

| Variable | Change in $R^2$ with other variables |
| --- | --- |
| Anti-Spike End point titre | 0.2277 |
| anti-Spike-IgM | 0.0519 |
| anti-RBD-IgG1 | 0.0456 |
| Disease severity (Severe) | 0.0309 |
| Days post symptom onset | 0.0297 |

| Variable | Change in R<sup>2</sup> with other variables |
| --- | --- |
| anti -RBD End point titre | 0.0235 |
| anti -RBD-IgM | 0.0235 |
| anti-Spike-IgG1 | 0.0234 |
| Neutralisation titre | 0.0189 |
| anti -RBD-IgG2 | 0.0159 |
| anti-Spike-IgG3 | 0.0127 |
| anti -RBD-IgA | 0.0115 |
| Age, years | 0.0092 |
| anti-Spike-IgG2 | 0.0027 |
| anti-Spike-IgG4 | 0.002 |
| anti -RBD-IgG3 | 0.0013 |
| anti -RBD-IgG4 | 0.001 |
| anti-Spike-IgA | 0.0002 |
| Gender (Female) | 0 |

**Fig 4:**
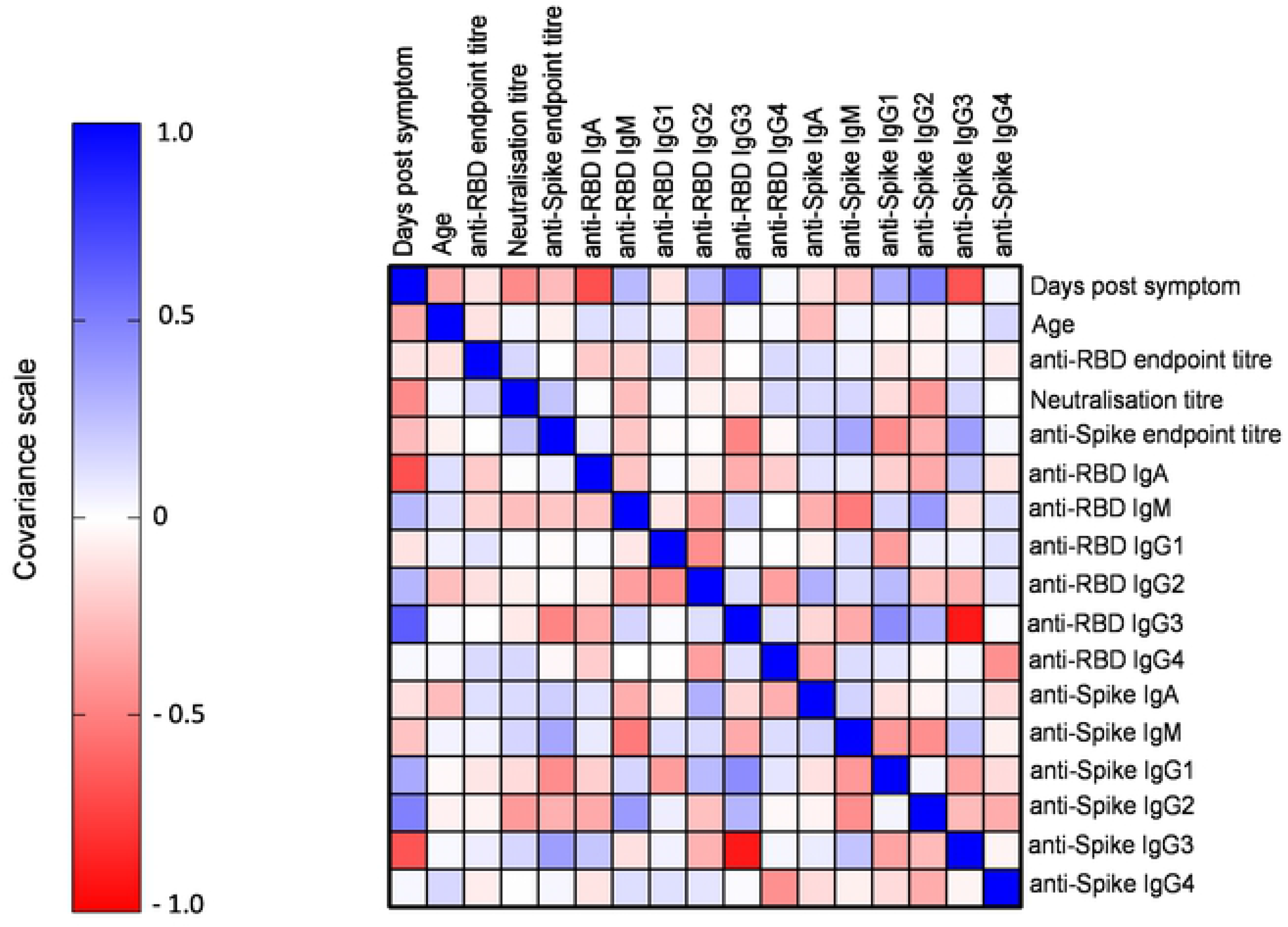
Spearman’s covariance matrix. A parameter Spearman’s covariance matrix to plots the associative relationship between the independent variable using the standardized beta coefficient covariance scale (range −1 to +1), where the score above one is considered a positive relationship, zero as no relationship and less than one as negative relationship.

### Anti-Spike p-score strongly correlates with endpoint antibody titre, neutralisation function and moderate disease

Spike p-scores significantly correlated with anti-Spike EPTs (Fig 5A, 5F) (Spearman r = 0.65, p = 0.0001), and neutralisation titre (Spearman r = 0.56, p = 0.0001) during convalescence (Fig 5B). Although neutralisation titre significantly correlated with anti-Spike EPT (Spearman’s r = 0.64, p = 0.0001) and anti-RBD EPT (Spearman’s r = 0.34, p = 0.009) (Fig 5C-D), it did not correlate with DPS. Stratification of patient group by disease severity indicated that patients with moderate disease had the most significant correlation of the Spike p-score to the anti-Spike EPTs (Spearman r = 0.71, p = 0.001), and to the neutralisation titre (Spearman r = 0.60, p=0.0007), while samples from individuals in the severe disease category had no significant correlation among the variables (Fig 5F).

**Fig 5:**
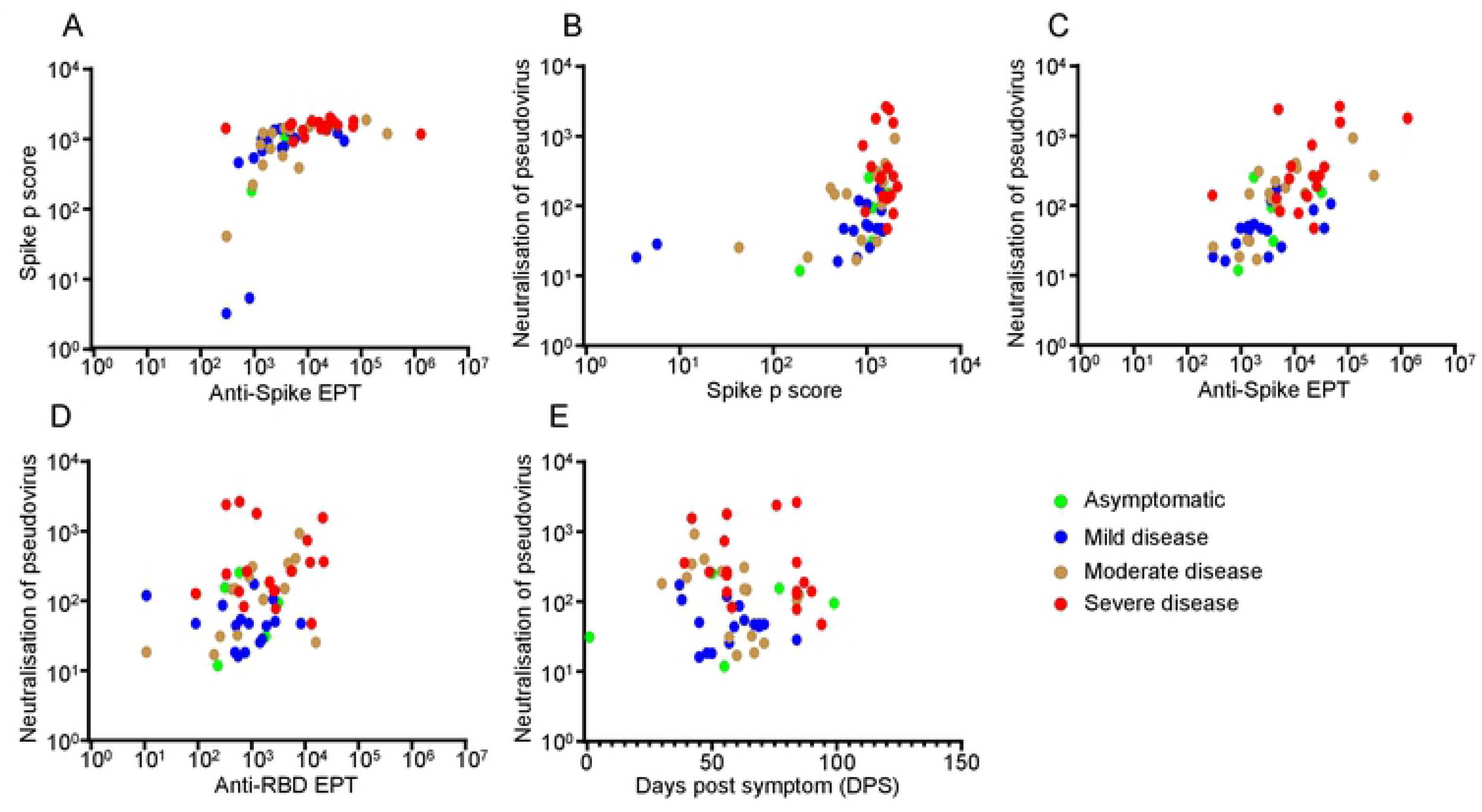
Correlation of Spike p-score to endpoint antibody titre and neutralisation titre in patients with varying disease severity. (A) Spearman correlation studies of all 56 convalescent patients showed significant positive correlation of Spike phagocytic scores with anti-Spike antibody EPTs and (B) the Spike pseudovirus neutralisation. (C) Stratification of the patients by disease severity indicated that patients with moderate disease had the most significant correlation of the Spike phagocytosis to the anti-Spike antibody EPTs and to the Spike pseudovirus neutralisation while patients at both ends of the spectrum had least significant correlation. Patients with moderate disease also displayed significant positive correlation of the neutralisation titre with the anti-Spike antibody EPTs (D) anti-RBD antibody EPTs and (E) days post symptom onset. (r and p values are shown in text and in supplementary Table 2).

Interestingly, patients with moderate disease also displayed a significant positive correlation between the neutralisation titre and the anti-Spike EPT (Spearman r = 0.7, p = 0.0005), as well as with anti-RBD EPT (Spearman’s r = 0.6, p = 0.01), while a negative but significant correlation with DPS was observed (Spearman’s r = −0.6, p = 0.007) (Figs 5C and 5F). There was no correlation between Spike p-score and anti-Spike EPTs in any of the age groups (Fig 6A). Only neutralisation titre correlated significantly with Spike p-score and anti-Spike EPT across the age groups (Fig 6B-C). Interestingly, only in the 40–60-year-old group, a significant correlation was observed between the neutralisation titre and anti-RBD EPT (Fig 6D).

**Fig 6:**
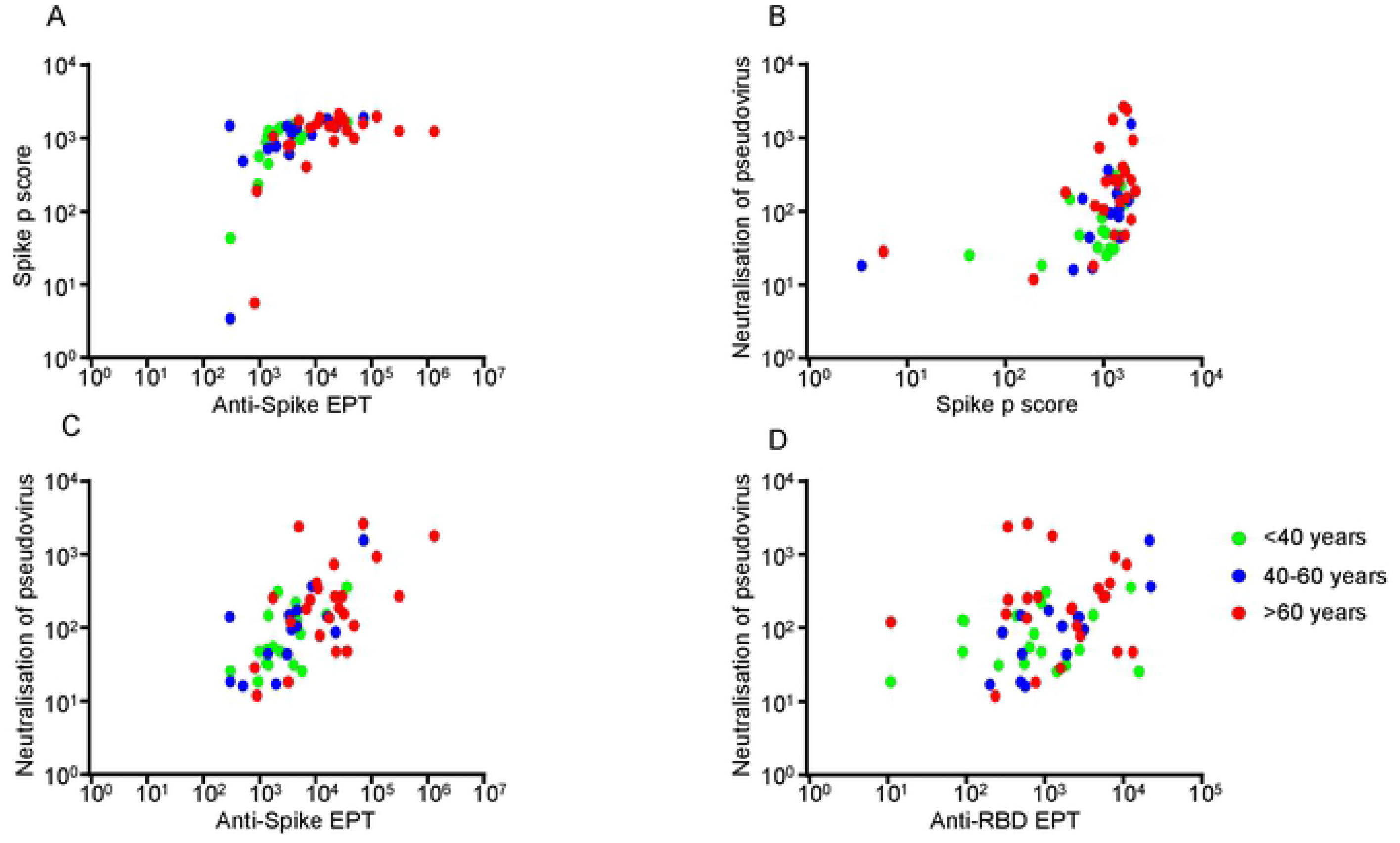
Correlation of Spike p-score to endpoint antibody titre and neutralisation titre in different age group patients. (A) Stratification of the convalescent patients by age into <40 years, 40-60 years and >60 years of age showed no significant correlation of the Spike phagocytosis to anti-Spike antibody EPTs in any of the age groups. (B) However, there was significant but variable correlation of the Spike phagocytosis to the Spike neutralisation titre in each age group. (C) All age groups showed significant correlation of their neutralisation titre to the anti-Spike EPTs. (D) Only the 40-60-year-old patients showed significant correlation of the neutralisation titre to anti-RBD antibody EPTs (r and p values are shown in text and supplementary Table 2).

### Maintenance of anti-Spike phagocytosis and neutralisation functions and increase in affinity of the anti-Spike antibodies despite decline of the endpoint antibody titers from baseline over 12-months follow-up

Further longitudinal study of nine convalescent patients showed a significant decline of anti-Spike IgG EPT [χ^2^ [2] = 18, p = 0.0001], Spike p-score [χ^2^ [2] = 12.6, p = 0.0007], and neutralisation function [χ^2^ [2] = 8, p = 0.01] from visit 1 (one month post symptom onset) to visit 3 (12 months post symptom onset)] regardless of disease severity (Fig 7A-C). The anti-Spike EPT significantly decreased from visit 1 to visit 3 (Friedmann test, visit 1-visit 3, p = 0.0001) (Fig 7A). Similarly, the Spike p-score significantly decreased from visit 1 to visit 3 (Friedmann test, visit 1-visit 3, p = 0.001) (Fig 7B). The neutralisation titre also decreased significantly from visit 1 to visit 3 (Friedmann test, visit 1-visit 3, p = 0.01) (Fig 7C). Although the anti-Spike EPT declined by an average of 91.2% (standard error of mean (SEM): 1.43) from visit 1 to visit 3, there was better maintenance of the Spike p-score and neutralisation titre, which showed 65.3% ± 9.27 SEM and 41.8% ± 39.2 SEM decline on average, respectively (Fig 7D). Interestingly, the surface plasmon resonance studies showed up to 840-fold increase in the affinity (K_D_) of the anti-Spike antibodies to Spike protein in 8/9 patient plasma samples (mean = 150-fold increase; range 3.3-836.8) (Fig 7E-F). Conversely, the binding avidity (K_d_) showed a significant decline in 8/9 patients by 115.8-fold (range 5.1-515.8) (Supplementary Fig 3A-B). These results suggest that an improvement in the quality of the antibodies over time may have contributed to retention of the phagocytic and neutralisation functions, despite the substantial decrease in the EPTs.

**Fig 7:**
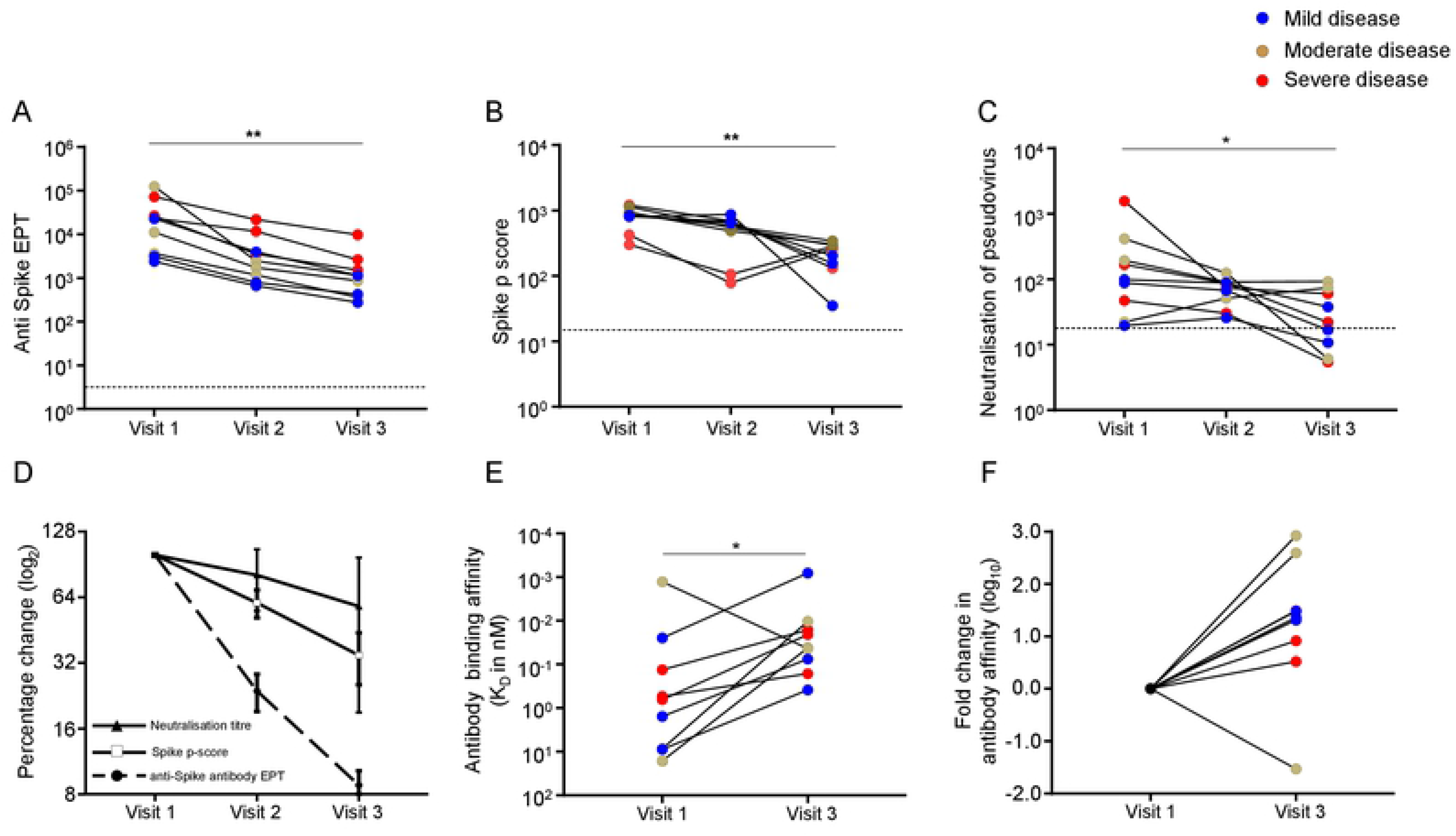
Longitudinal study of anti-Spike antibodies EPTs, Spike phagocytosis, Spike neutralisation titre and affinity of plasma anti-Spike antibodies. Longitudinal study of nine convalescent patients with mild, moderate, or severe disease (n=3 each) showing significant time-dependent decline of (A) anti-Spike IgG antibodies endpoint titers, (B) Spike phagocytosis, and (C) anti-Spike neutralisation titre over a 12-months period regardless of their disease severity. (D) The anti-Spike EPT declined by average 91.2% (standard error of mean (SEM): 1.43) from visit 1 to visit 3, but there was substantial retention of the Spike p-score and neutralisation titre, which showed 65.3% ± 9.27 SEM and 41.8% ± 39.2 SEM decline on average, respectively. (E-F) Surface plasmon resonance showing increase in the affinity (K_D_) of the anti-Spike antibodies in patient plasma to recombinant Spike protein in 8/9 patients between visit 1 and visit 3 by up to 840-fold (mean = 150-fold increase; range 3.3-836.8). Dotted lines in each plot show median or mean values +3SD of 25 healthy control samples (*p<0.05, **p<0.01).

## Materials and Methods

### Study cohort

Stored frozen plasma samples from 62 patients enrolled to an ongoing prospective cohort study (COSIN-Collection of Coronavirus COVID-19 Outbreak Samples in New South Wales-2020) [20] and from Central Adelaide Health Network (CALHN) were used in this study. Plasma was collected on days 10-94 post symptom onset. In a subset of patients (n=9), two additional follow up samples were collected on days 110-252 and days 321-381 post symptom onset. This subset was chosen based on both availability of samples and representation of the whole spectrum of disease severity. Infection with SARS-CoV-2 was confirmed in all patients by a validated quantitative RT-PCR used by diagnostic laboratories across NSW. Disease severity was classified according to the NIH COVID-19 treatment guidelines [21]. Archival serum or plasma collected from 25 healthy donors prior to the pandemic with an age range of 24-73 years, and a male to female ratio of 1:2.4 was used as controls. Buffy coats from 3 healthy donors obtained from the Australian Red Cross (material transfer agreement # 18-01NSW-06) were used as a source of blood monocytes for *in vitro* differentiation of primary macrophages. The control archival plasma samples were collected prior to 2019, and the serum as well the buffy coats were collected before April 2020 when local transmission was low in NSW and none of the donors were close contacts of patients with COVID-19.

The study protocol was approved by the Human Research Ethics Committees of the Northern Sydney Local Health District, the University of New South Wales, NSW Australia (ETH00520), CALHN Human Research Ethics Committee, Adelaide, Australia (Approval No. 13050), and the Women’s and Children’s Health Network Human Research Ethics (protocol HREC/19/WCHN/65), Adelaide, Australia which was conducted according to the Declaration of Helsinki and International Conference on Harmonization Good Clinical Practice (ICH/GCP) guidelines and local regulatory requirements. Written informed consent was obtained from all participants before enrolment.

### Cells and antibodies

The monocytic cell line, THP-1 was obtained from ATCC 202 TIB [22]. THP-1 cells were cultured in RPMI 1640 supplemented with 2mM L-glutamine (Gibco, USA),10% Fetal bovine serum, 0.05mM β-mercaptoethanol, 10mM HEPES and 100U/mL Penicillin-Streptomycin (Thermofisher Scientific, USA) and passaged every two days. Expression of surface Fc-receptors was assessed by flow cytometry using monoclonal antibodies (mAbs) against Fcγ receptor RI (CD64)-FITC, FcγRIII (CD16)-PE, CD14-PerCP (Becton Dickinson, USA) and FcγRII (CD32)-ACP (Life technologies, USA) and isotype and fluorochrome matched negative control mAbs (BD) and used for the phagocytosis at passages 5-10 [23].

Peripheral blood mononuclear cells were isolated from buffy coats by gradient centrifugation (Lymphoprep Stem cell, USA), resuspended in RPMI-1640 complete media containing 10% human AB serum (Sigma) at 2×10^6^/mL and seeded onto 24 well flat bottom plates containing poly-L-Lysine (Sigma, USA) coated glass cover slip inserts (Deckglasser, Germany). After a one hour incubation in a humidified 37°C incubator with 5% CO_2_ air, non-adherent cells were removed by washing wells twice with pre-warmed PBS and the adherent monocytes (~1×10^4^/well) were differentiated into macrophages for six days using 50ng/mL granulocyte-macrophage colony-stimulating factor and 25ng/mL interleukin-10 (Sigma, USA), as described previously [24].

### Biotinylated recombinant SARS-CoV-2 Spike and RBD proteins

SARS-CoV-2 Wuhan-Hu-1 (GenPept: QJE37812) RBD protein (amino acid residues 319–541) and Spike protein (amino acid residue 15-1213) were cloned into pCEP4 mammalian expression vector containing N-terminal human Ig kappa leader sequence and C-terminal Avi-tag and His-tag (Invitrogen, USA). Expi293-Freestyle cells cultured at 37°C and 8% CO_2_ in growth medium containing Expi293 Expression Medium at 3×10^6^/mL in 50 mL media were transfected overnight at 37 °C with 50μg of plasmid in 160μL of ExpiFectamine plus 6mL of OptiMEM-I (all from Invitrogen). The following day 300μL of ExpiFectamine Enhancer 1 and 3mL of Enhancer 2 (Invitrogen) was added and the secreted recombinant Spike or RBD protein in culture supernatant were harvested after 72 hours and affinity purified using HisTrap HP Column (GE Healthcare, USA) as described previously [25]. The purified Spike and RBD recombinant proteins were then buffer exchanged into sterile PBS by centrifuging at 4000xg for 30 minutes at 4 °C in a 10,000 MWCO Vivaspin centrifugal concentrator (Sartorius, Germany) and stored at –80 °C. The Spike and RBD recombinant proteins were then biotinylated via the C-terminal Avi-tag using a labelling kit following the manufacturer’s instructions (Genecopeia, USA) [20,26].

### Endpoint titre and Isotyping of anti-SARS-CoV-2 Spike and anti-RBD antibodies in sera of patients with SARS-CoV-2 infection

Anti-SARS-CoV-2 Spike and RBD IgG antibody in sera was quantified using a modified direct ELISA [27]. Briefly, Nunc immuno-microtiter plates were coated with 250ng of the recombinant SARS-CoV-2 RBD or 100ng of SARS-CoV-2 Spike protein per well for 2 hours at room temperature and washed three times with 137mM NaCl, 2.7mM KCl, 8mM Na_2_HPO_4_, and 2mM KH_2_PO_4_, 0.1% Tween 20 (wash buffer) to remove unbound protein. After blocking for one hour at room temperature with 5% skim milk in wash buffer, plates were washed once, and serially diluted serum (in 5% skim milk) was added to each well in duplicate, incubated for one hour at room temperature and plates washed twice with wash buffer. For the endpoint titre of total anti-Spike or anti-RBD antibodies, 50μL of horseradish peroxidase (HRP)-conjugated mouse anti-human detection antibody added per well (Jackson ImmunoResearch, USA) (1:5000 dilution in 5% skim milk). For isotyping of the anti-Spike or anti-RBD antibodies, 50μL/well of HRP-conjugated immunoglobulin subtype or IgG subclass specific detection antibodies were added. These include 1:6000 dilution of anti-total human IgG (Jackson Immunoresearch), 1:3000 dilution of anti-human IgA (α-chain-specific) (Sigma) or 1:3000 dilution of anti-human IgM (μ-chain-specific) (Sigma) antibody subtypes and 1:6000 dilution of anti-human IgG1, IgG2, IgG3 or IgG4 (Southern Biotech, USA). After one hour incubation with the HRP-conjugated detection antibodies at room temperature, wells were washed twice with wash buffer, incubated with 3,3′,5,5′-Tetramethylbenzidine horse radish peroxidase substrate (50μL per well) for 10 minutes, the colorimetric reaction stopped by adding 50μL/well of 1M HCl (Sigma, USA) and optical density at 450 nm measured using CLARIOstar microplate reader (BMG Labtech, Australia).

### SARS-CoV-2 Spike or RBD protein coating and opsonisation of microbeads

1.5×10^9^beads/mL streptavidin and Alexa-488 florescent tagged 0.4 μm polyester microbeads (Sphereotech, USA) were mixed with recombinant biotinylated SARS-CoV-2 Spike or RBD protein (at 50μg/mL) in 1.5mL Eppendorf tubes and incubated for 16 hours at 4°C on a rotating chamber. Excess protein was removed by washing the beads twice with 1mL lipopolysaccharide minimised cold PBS and gentle centrifugation at 2292xg for five minutes (Beckman coulter microfuge 20R). Aliquots (50μL) of the SARS-CoV-2 Spike or RBD coated beads were then opsonised for two hours at 37°C with 10μL of plasma obtained from patients with SARS-CoV-2 infection, or from healthy control donors, and used immediately in the antibody dependent cellular phagocytosis assay.

### SARS-CoV-2 specific phagocytosis assays

THP-1 cells in PBS (1×10^5^ cells in 50μL) were added onto 60μL of the patient plasma opsonised microbeads in a 1.5mL Eppendorf tube, mixed by gentle tapping, adjusted to 600μL using RPMI 1640 containing 0.1% human serum and 100mM HEPES and transferred into 37°C, 5% CO_2_ incubator. After a two hours incubation, cells were washed once with 1mL of cold PBS containing 0.5% FBS and 0.005% of sodium azide and gentle centrifugation at 335xg for five minutes at 4°C, fixed in 400μL of 1% paraformaldehyde and kept at 4°C in the dark until acquisition of data using BD FACSCalibur™ Flow cytometer. A total of 2×10^4^ events were acquired and the proportions of cells that phagocytosed the beads (% of cells that took up the beads) and their fluorescent intensities (amounts of beads taken up per cell) were analysed using BD FlowJo version 10.5.0 software. Phagocytic scores (p-score) were then calculated based on the proportion of cells that took up the opsonised beads, and mean fluorescence intensity (MFI) representing the average bead uptake by the positive cells, as described [23]. Cells incubated with Spike- or RBD protein-coated microbeads opsonised with plasma from healthy donors were used as negative controls. A positive p-score was defined as three standard deviations above the background mean phagocytic score of healthy donors, as described [23].

To assess whether the uptake of the microbeads opsonised with the plasma of patients with acute disease was via Fc-receptor (antibody-dependent), and other heat labile opsonins such as complements (eg C3b), either the Fc-receptors on the THP-1 cells were pre-blocked using the universal Fc receptor blocking agent (Miltenyi, USA) [23], or the plasma heat inactivated at 56°C for 30 minutes, as described [28].

In selected experiments, the intracellular uptake of the opsonised microbeads by effector cells was confirmed by confocal microscopy as described [23]. In brief, primary macrophages (1×10^4^ cells) on poly-L-Lysine (Sigma, USA) coated glass cover slips were rinsed with PBS, resuspended in RPMI 1640 containing 0.1% human serum and 100mM HEPES and incubated with the Alexa-488 conjugated opsonised microbeads for two hours at 37°C, 5% CO_2_. Cells were then washed twice with cold PBS containing 0.5% FBS and 0.005% sodium azide, fixed with 1% paraformaldehyde for five minutes at room temperature, and rinsed twice with PBS. The fixed cells were blocked with 1% BSA in PBS, incubated with 1:1000 dilution of Alexa-555-conjugated Phalloidin (Sigma, USA) for 30 minutes at room temperature, mounted in DAPI nuclear stain containing media (Molecular Probes, USA) and imaged using ZEISS LSM 880 confocal microscope (Carl Zeiss AG, Germany), using 63X/1.4 Plan-Apochromat Oil Immersion objective, with Diode 405 nm (DAPI), Argon ion 488 nm (Alexa-488) and DPSS 561 nm (Alexa-555 phalloidin) laser excitation sources, emitted light was filtered using combination of emission filters and imaged onto Airy detector array producing an effective lateral resolution of ~100 nm. All the images were Airyscan processed with Zen Black Edition (Zeiss Software).

### SARS-CoV-2 pseudovirus neutralisation assay

Retroviral SARS-CoV-2 Spike pseudovirus were generated in 293T cells by co-transfecting expression plasmids containing SARS-CoV-2 Spike and MLV gag/pol and luciferase vectors using Calphos transfection kit (Takara Bio, USA) as described [20]. Pseudovirus in culture supernatants were then harvested 48 hours post transfection, concentrated 10-fold using 100,000 MWCO Vivaspin centrifugal concentrators (Sartorius, Germany) and used for neutralisation assays. Briefly, pseudovirus were incubated for one hour with heat inactivated (56 °C for 30 minutes) patient serum prior to infecting 293T-ACE2 cells (kindly provided by A/Prof Jesse Bloom) by a two hour spinoculated at 800xg in 96-well white flat bottom plates in triplicates (Sigma-Aldrich, USA). Infected cells were incubated at 37°C in a humidified incubator with 5% CO_2_, replenished with fresh media, further incubated for 72 hours, and then lysed with a lysis buffer (Promega, USA). Relative Luminescence Unit (RLU) in cell lysates was measured using CLARIOstar microplate reader (BMG Labtech, Australia), percentage neutralisation of Spike pseudovirus determined and the fifty percent inhibitory (ID50) calculated using non-linear regression model (GraphPad Prism version 9.0) [29]. Positive ID50 cutoff was defined as two standard deviations above the mean background reading obtained from 10 healthy subjects.

### Surface plasmon resonance

Surface plasmon resonance (SPR) was performed using a Biacore T200 (Cytiva, USA) to determine the binding characteristics of the anti-Spike polyclonal antibodies in patient plasma to SARS-CoV-2 Spike antigen. Briefly, 10μg/mL of C-terminal 6X-His tag containing recombinant SARS-CoV-2 Spike protein was captured at a flow rate of 10μL/minute for 180 seconds onto carboxymethylated (CM5) dextran sensor chips immobilised with anti-His mAb using EDC/NHS amine coupling kit (GE Healthcare, Australia). The sensor chips were equilibrated with running buffer [HEPES buffer (HBS-EP+; 0.01M HEPES pH7.4, 0.15M NaCl, 3mM EDTA, 0.05% v/v Surfactant P20] before the addition of plasma. Plasma from patients with SARS-CoV-2 infection or healthy controls diluted (1:100) in the running buffer were then injected over the immobilised flow cells at a rate of 20 μl/min for 120 s at 25°C. After each patient plasma, then chip was regenerated using 10 mM glycine, pH 2 (Cytiva, USA) for 60 seconds. Binding responses with plasma were double reference-subtracted from non-specific responses to an empty flow cell and blank injection (zero analyte concentration). Kinetic constants, including association constant (*K*_a_), equilibrium constant (*KD*) (affinity) and dissociation constant (*Ka*) (avidity), were calculated using BIAcore evaluation software version 4.1 [30].

### Statistical analysis

All data were analysed with Prism Software (version 9.0, GraphPad, USA). Unpaired non-parametric Mann Whitney U-tests were used to compare p-scores (Spike/RBD), neutralisation titre, antibody end point titre (Spike/RBD) between patient groups/subgroups and healthy controls. To test the difference between three or more groups, parametric ANOVA or Kruskal-Wallis test followed by Dunn’s test was used, where appropriate, based on the distribution of the residual plot. Spearman’s correlation was used to compare Spike and RBD end point titre against Spike p-score/RBD p-score/neutralising titers. Friedman’s test with pairwise Dunn’s tests were used to compare the repeated measures of Spike p-score, Spike and RBD end point titre, neutralisation titre and antibody affinity across the timepoints V1, V2 and V3.

To rank the variables of statistical importance that associated with Spike p-score (dependent variable), multiple linear regression analysis was performed using disease severity, age, gender, DPS, anti-Spike/RBD endpoint antibody titre, anti-Spike/RBD antibody subtypes and neutralisation titre as independent variables. The variables were ranked based on the change in R^2^ and by treating each variable as the last one that entered the regression model. This change in R^2^ represents the amount of unique variance that each variable explains that the other variables in the model cannot explain. A parameter covariance matrix was then used to plot the associative relationships between the independent variable using the standardised beta coefficient, where a score above one is considered a positive relationship, zero as no relationship and less than one as a negative relationship.

## Discussion

The studies reported here demonstrated that acutely ill patients with COVD-19 can mount phagocytic responses mediated by heat labile components of the plasma as early as 10 days post symptom onset, regardless of disease severity and anti-Spike end point antibody titers. COVID-19 disease severity was also found to be associated with increased phagocytic responses in patients that recovered from COVID-19 disease. Importantly also, this report reveals for the first time that affinity of anti-Spike antibodies in convalescent patient samples increased over the 12-month period leading to retention of the phagocytic and neutralisation functions, despite a marked decline in the antibody titers.

Studies to date have used antibodies purified from patient samples, or cloned monoclonal antibodies, to detect phagocytosis, revealing a narrow pattern of antibody dependent phagocytosis without recognition of significant contribution of plasma proteins such as activated complement products that are recognised to be critical in phagocytic responses against other respiratory pathogens [11,17,18]. By contrast, in this study, native patient plasma was used revealing that the early phagocytic response in SARS-CoV-2 infection is primarily driven by heat labile components in the plasma, along with a contribution from Fc-receptor dependent antibody functions. Further, the studies revealed that samples from acutely ill patients may have significant anti-Spike antibody mediated phagocytosis activity, despite having undetectable antibody titers. These findings suggest that like other respiratory viral infections, the most important heat labile components of plasma contributing to viral phagocytosis are the classically activated complement products that upon interaction with complement-fixing IgM and IgG1 can lead to enhanced phagocytosis through interactions with Fc and complement receptors [9,31], or by direct opsonisation of viruses by complement proteins activated via the mannose-binding lectin pathway [11–14,17].

There is ample evidence to support the association between complement activation through the classical, lectin and alternate pathways and clinical severity in COVID-19, including strong links to worsened disease severity and systemic inflammation [11–14]. Interestingly, it has also been shown that anti-Spike antibodies from patients with acute COVID-19 can initiate tissue deposition of complement that may initiate tissue injury [32]. Our finding that blocking of the Fc-receptors can only partially abrogate the phagocytic response, supports the notion of collaborative effects between complement proteins and the Fc-receptor specific antibody functions. Enhancement of phagocytosis by complement may trigger Fc-receptor- and complement receptor-mediated excessive activation of phagocytic effector cells leading to the hyperinflammatory state likely underpinning the severe disease observed in some members of the cohort. This collaborative response may also contribute to more effective viral elimination leading to the sometimes favourable clinical outcome, as observed by Federica et al [11].

Although the observations in acutely ill patients are potentially clinically significant, the limited numbers available from this early stage of the disease preclude definitive conclusions, hence further studies in a larger cohort are warranted. By contrast, in the convalescent patients phagocytosis strongly correlated with anti-Spike antibody endpoint titers, neutralisation functions and severe disease. Multiple regression analysis showed that the most important variable related to a high phagocytic response was anti-Spike antibody endpoint titre. Further, older patients mounted significantly higher neutralisation activity and had higher antibody endpoint titers, while gender had little impact on the phagocytic function. These results are consistent with recent studies in which disease severity and age were identified as major contributors to endpoint antibody titers and their neutralisation functions [32–34]. While co-morbidities may have contributed to disease severity in the older patients studied here, it is also possible that pre-existing cross-reactive antibodies against other common coronaviruses may have led to consistently higher anti-Spike antibody titers [6], that in turn may explain the higher phagocytic responses. Future, longitudinal characterisation of phagocytic and neutralisation functions of anti-Spike antibodies against emerging variants of SARS-CoV-2 infection and their functional cross-reactivity to the earlier variants are warranted.

To date, phagocytic activity in SARS-CoV-2 has been reported to be maintained for 5 months post infection [16]. The findings in the longitudinal study reported here indicated retention of both phagocytic and neutralisation functions for over 12 months, despite a progressive decline in anti-Spike antibody endpoint titers. While the progressive decline of the antibody titers is consistent with previous studies [35–37], the data presented here show for the first time, a significant increase in the anti-Spike antibody affinity over time, which may have contributed to the retention of the effector functions despite the decay of the endpoint titers to baseline. This finding concurs with longitudinal B cell studies that have reported ongoing affinity maturation and improved affinity of SARS-Cov-2 specific memory B cell receptors up to 12 months [38]. Importantly, this report is the first study to demonstrate antibodies in the circulation showing functional improvement correlating with the functional characteristics observed in memory B cells. These new results have potential clinical implications. They challenge current clinical practice that considers measurement of the anti-Spike antibody titers but not their quality as one of the gold standard indicators of immune protection after SARS-CoV-2 infection or vaccination. Change in practice that includes routine measurement of the quality and functions of these antibodies is therefore highly recommended.

## Acknowledgements

The authors would like to thank the study participants for their contribution to the research, as well as current and past researchers and staff. They would like to acknowledge members of the study group: Protocol Steering Committee – Rowena Bull (Co-Chair, The Kirby Institute, UNSW Sydney, Sydney, Australia), Marianne Martinello (Co-Chair, The Kirby Institute, UNSW Sydney, Sydney, Australia), Andrew Lloyd (The Kirby Institute, UNSW Sydney, Sydney, Australia), John Kaldor (The Kirby Institute, UNSW Sydney, Sydney, Australia), Greg Dore (The Kirby Institute, UNSW Sydney, Sydney, Australia), Tania Sorrell (Marie Bashir Institute, University of Sydney, Sydney, Australia), William Rawlinson (NSWHP), Jeffrey Post (POWH), Bernard Hudson (RNSH), Dominic Dwyer (NSWHP), Adam Bartlett (SCH), Sarah Sasson (UNSW) Nick Di Girolamo (UNSW) and Daniel Lemberg (SCHN).

Coordinating Centre - The Kirby Institute, UNSW Sydney, Sydney, Australia – Rowena Bull (Co-ordinating principal investigator), Marianne Martinello (Co-ordinating principal investigator), Marianne Byrne (Clinical Trials Manager), Mohammed Hammoud (Post-Doctoral Fellow and Data Manager), Andrew Lloyd (Investigator) and Roshana Sultan (Study co-ordinator).

Site Principal Investigators – Jeffrey Post (Prince of Wales Hospital, Sydney, Australia), Michael Mina (Northern Beaches, Sydney, Australia), Bernard Hudson (Royal North Shore Hospital, Sydney, Australia), Nicky Gilroy (Westmead Hospital, Sydney, Australia), William Rawlinson (New South Wales Health Pathology, NSW, Australia), Pam Konecny (St George Hospital, Sydney, Australia), Marianne Martinello (Blacktown Hospital), Adam Bartlett (Sydney Children’s Hospital, Sydney, Australia) and Gail Matthews (St Vincent’s Hospital, Sydney, Australia).

Site co-ordinators – Dmitrii Shek and Susan Holdaway (Blacktown hospital), Katerina Mitsakos (RNSH), Dianne How-Chow and Renier Lagunday (POWH), Sharon Robinson (SGH), Lenae Terrill (NBH), Neela Joshi, (Lucy) Ying Li and Satinder Gill (Westmead), Alison Sevehon (SVH).

---

**Supplementary Fig 1: Optimisation of SARS-CoV-2 specific phagocytosis assay using Flow cytometry**.

(A) Representative dot plots showing proportion of THP-1 effector cells phagocytosing Spike protein coated non-opsonised fluorescent microbeads as background control. (B) Spike protein coated fluorescent microbeads opsonised with healthy control plasma as negative control, and (C) Spike protein coated fluorescent microbeads opsonised with plasma of a COVID-19 patient with high anti-Spike antibody EPT as positive control; percentages of positive phagocytosis shown on the lower right and upper right quadrants of each plot. (D) Representative histograms showing the mean fluorescence intensity (MFI) of the THP-1 cells that phagocytosed the fluorescent microbeads; green = Spike protein coated non-opsonised fluorescent microbeads as background control; red = Spike protein coated fluorescent microbeads opsonised with healthy control plasma and blue = Spike protein coated fluorescent microbeads opsonised with a plasma of a COVID-19 patient. (E) Representative confocal microscopic images of peripheral blood derived primary macrophages showing minimal uptake of the Spike protein coated fluorescent microbeads opsonised with healthy control plasma (upper panel) as contrasted to the extensive uptake of Spike protein coated fluorescent microbeads opsonised with a plasma of a COVID-19 patient (lower panel).

**Supplementary Fig 2: Sex dependent differences in endpoint antibody titers, phagocytosis and neutralisation titre at convalescence**.

(A) There was no significant difference in either anti-Spike antibody end point titre or (B) anti-RBD antibody end point titre; or (C, left) phagocytic responses between male and female patients; although C, right) there was significantly higher anti-Spike neutralisation titre in males.

**Supplementary Fig 3: Longitudinal assessment of binding avidity of anti-Spike antibodies in patient plasma**.

(A) Surface plasmon resonance studies using plasma from convalescent patients showed significant decline in the binding avidity (K_d_) of their anti-Spike antibodies to recombinant Spike protein in 8/9 patients at the 12-month time point (visit 3) compared to visit 1 (30-40 days post symptom onset) (*p<0.05); (B) with an average decline of 115.8-fold (range 5.1-515.8).

